# microRNAs Over-expressed in Diabetic Foot Ulcers Healing - Computational Modeling of Molecular Structure

**DOI:** 10.1101/2021.07.16.452675

**Authors:** Luis Jesuino de Oliveira Andrade, Alcina Maria Vinhaes Bittencourt, Luís Matos de Oliveira, Gabriela Correia Matos de Oliveira

## Abstract

**Background:** Vasculopathy associated with diabetic neuropathy are important risk factors for the diabetic foot ulcers development. Diabetic foot ulcers is severe complication that occur in about 15% of people with diabetes, being able require hospitalization and amputation in its treatment.

**Objective:** Design *in silico* the molecular structure of micro-ribonucleic acid (miRNA) overexpressed in diabetic foot ulcers healing.

**Method:** We performed a careful search of the nucleotide sequence of 8 miRNAs over-expressed in diabetic foot ulcers, designing in silico the molecular structure of following miRNAs: miRNA-146a, miRNA-155, miRNA-132, miRNA-191, miRNA-21, miRNA-203a, miRNA-203b, and miRNA-210. The nucleotides were taken from GenBank of National Center for Biotechnology Information genetic sequence database. The sequences acquired were aligned with the Clustal W multiple alignment algorithms. The molecular modeling of structures was built using the RNAstructure, an automated miRNAs structure modelling server.

**Results:** We showed a search for nucleotide sequence and the design of the molecular structure of following miRNA over-expressed in diabetic foot ulcers healing: miRNA-146a, miRNA-155, miRNA-132, miRNA-191, miRNA-21, miRNA-203a, miRNA-203b, and miRNA-210. We produced a tutorial on a molecular model of the 8 miRNAs overexpressed in the diabetic foot by processing *in silico* projection of their molecular structures.

**Conclusion:** We show *in silico* secondary structures design of selected of 8 miRNAs over-expressed in diabetic foot ulcers healing by means of computational biology.

## INTRODUCTION

Diabetic foot is a serious diabetic complication, associated to peripheral vascular disease and diabetic neuropathy, resultant of ulcerations in deep tissues in the lower limbs.^1^ Epidemiological studies show that the diabetic foot is a complication that affects about 15% of diabetic patients and that 85% of all amputations due to diabetic foot are preceded by a foot ulceration that may evolve with infection or gangrene.^2^ The diabetic foot has a pathophysiological basis in neuropathy, trauma, and sometimes associated with occlusive arterial disease of the lower limbs, leading to the risk of ulceration with soft tissue infection.^3^ Micro-ribonucleic acid (MiRNAs) are controlling molecules that are associated with various expressions and complications of diabetes and several microRNAs have been linked to the progression and severity of diabetic foot.^4^

MiRNAs are tiny non-coding RNAs composed of 19 to 35 nucleotides involved in the post-transcriptional control of gene expression in multicellular organisms.^5^

In recent years the understanding about miRNA structure and function has increased expressively. The bioinformatics software available for creating molecular models and investigating nucleotide sequences provide the necessary devices for assembling and understanding the molecular mechanisms of miRNAs. The aim of this study was to design *in silico* the molecular structure of 8 miRNA overexpressed in diabetic foot ulcers, and to produce a tutorial on its modeling.

## METHODS

We performed a careful search of the nucleotide sequence of 8 miRNAs over-expressed in diabetic foot ulcers, designing in silico the molecular structure of following miRNAs: miRNA-146a, miRNA-155, miRNA-132, miRNA-191, miRNA-21, miRNA-203a, miRNA-203b, and miRNA-210.

The nucleotides were taken from GenBank of National Center for Biotechnology Information genetic sequence database (https://www.ncbi.nlm.nih.gov/). The sequences acquired were aligned with the Clustal W multiple alignment algorithms. The molecular modeling of structures was built using the RNAstructure, an automated miRNAs structure modelling server (http://rna.urmc.rochester.edu/RNAstructureWeb/).

We produced a tutorial on a molecular model of the 8 miRNAs overexpressed in the diabetic foot by processing in silico projection of their molecular structures.

### Nucleotide search and sequence analysis

GenBank is a public domain nucleotide sequence analysis application available at https://www.ncbi.nlm.nih.gov/genbank/submit/. It has multiple nucleotide search algorithms and a wide variety of nucleotide algorithms are used to search many different sequence databases. In addition GenBank has the Nucleotide, Genome Survey Sequence (GSS) and the Expressed Sequence Tag (EST) that contain nucleic acid sequences. The information in EST and GSS comes from sharing sequencing quantities from GenBank.

### Molecular model building

The function and conformation of proteins and amino acids are established by nucleotide sequences and structure prediction still remains a significant difficulty, with an immense demand for high-resolution structure assumption methods.

Homology modeling is currently the most accurate computational method for generating reliable structural models and is commonly employed in multiple biological applications.

### Modeling the structure of RNA

The RNAstructure server determines an unfolding function, meets structures with max predictable precision, defines the maximum free energy structure, and pseudoknot predicts. In addition, this server generates sets of secondary structures with comments on highly possible correlations, starts with low free energy structure and includes others with variable correction expectations. If the minimum free energy structure is not appropriate several structures may be incorporated. Moreover, a second group of structures will be produced with shape constraints. The RNAstructure package is free of charge available for public use and download, from the Mathews lab homepage at: http://rna.urmc.rochester.edu/RNAstructureWeb.

## RESULTS

We showed a search for nucleotide sequence and the design of the molecular structure of following miRNA over-expressed in diabetic foot ulcers healing: miRNA-146a, miRNA-155, miRNA-132, miRNA-191, miRNA-21, miRNA-203a, miRNA-203b, and miRNA-210. We produced a tutorial on a molecular model of the 8 miRNAs overexpressed in the diabetic foot by processing *in silico* projection of their molecular structures.

### Nucleotide sequence of miRNA-146a

To construct the structure of miRNA-146a the nucleotide sequences with the NCBI identifier code: NR_029701.1 were used under the FASTA format obtained from GenBank. The miRNA-146a was planned to encode a 99 bp linear. Homo sapiens microRNA 146a (MIR146A) analysis is demonstrated in Figure 1.

**Figure 1.**
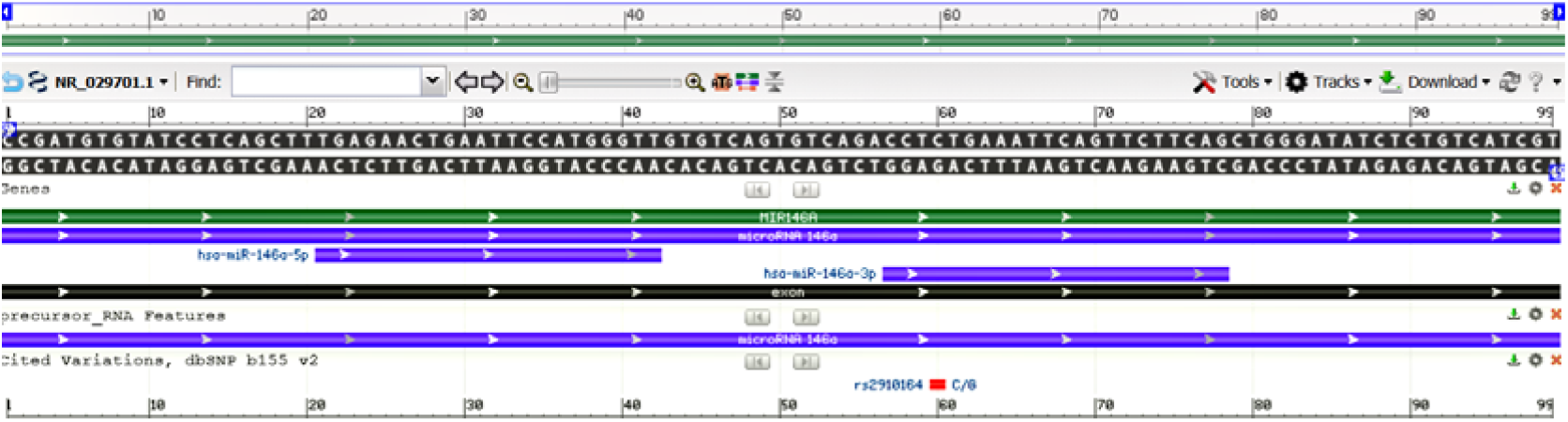
Homo sapiens microRNA 146a (MIR146A) - *model-template aligment*. **Source**: https://www.ncbi.nlm.nih.gov/nuccore/NR_029701.1?report=graph

### Molecular model of miRNA-146a

Nucleotide sequences of Homo sapiens microRNA 146a (MIR146A) were acquired employing FASTA format; modeling was performed employing the RNAstructure server, optimized and adjusted for alignment between structural templates and miRNA-146a nucleotide. Based on sequence alignment between the template structure and miRNA-101 nucleotide, a structural model was built for the nucleotide in question. So, employing RNAstructure server of comparative nucleotide modeling we generated a homology model of microRNA 146a (MIR146A), demonstrated in Figure 2.

**Figure 2.**
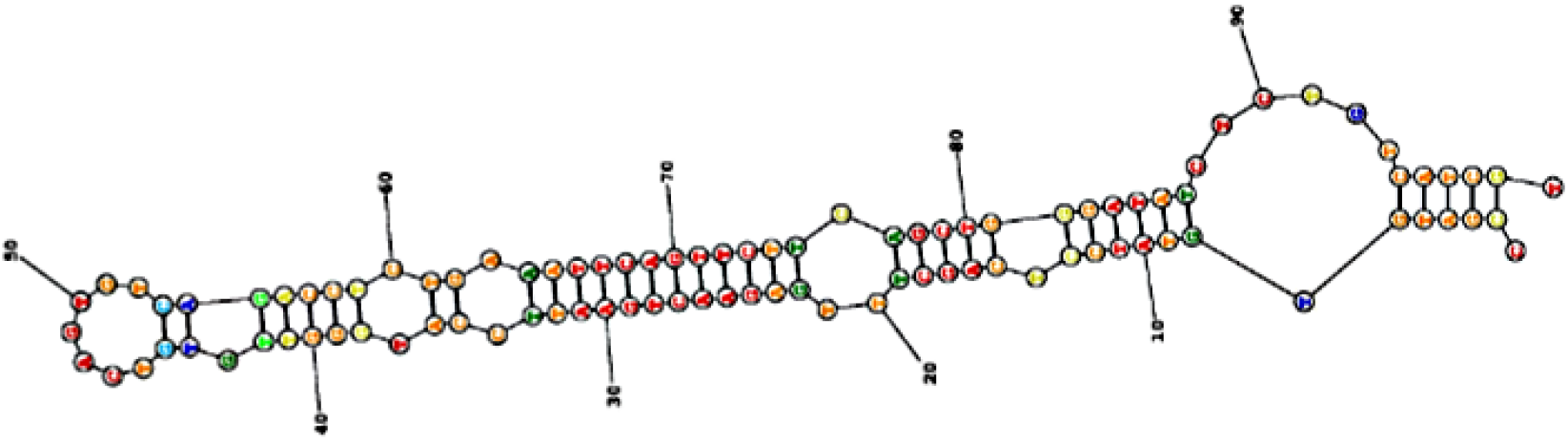
Homology model of the Homo sapiens microRNA 146a (MIR146A). **Source**: https://rna.urmc.rochester.edu/RNAstructureWeb/Servers/Predict1/ResultsPages/20210703.122549-6f09202a/Results.html

### Nucleotide sequence of miRNA-155

The reconstruction of miRNA-155 was performed from a nucleotide sequence archive in FASTA format obtained in GenBank database with the identifier code NCBI Reference Sequence: NR_030784.1. The miRNA-155 was planned to encode a 65 bp linear ncRNA. All coded sequences selected in FASTA format, used the annotation of the NCBI - Graphics. The Homo sapiens microRNA 155 (MIR155) analysis is shown in Figure 3.

**Figure 3.**
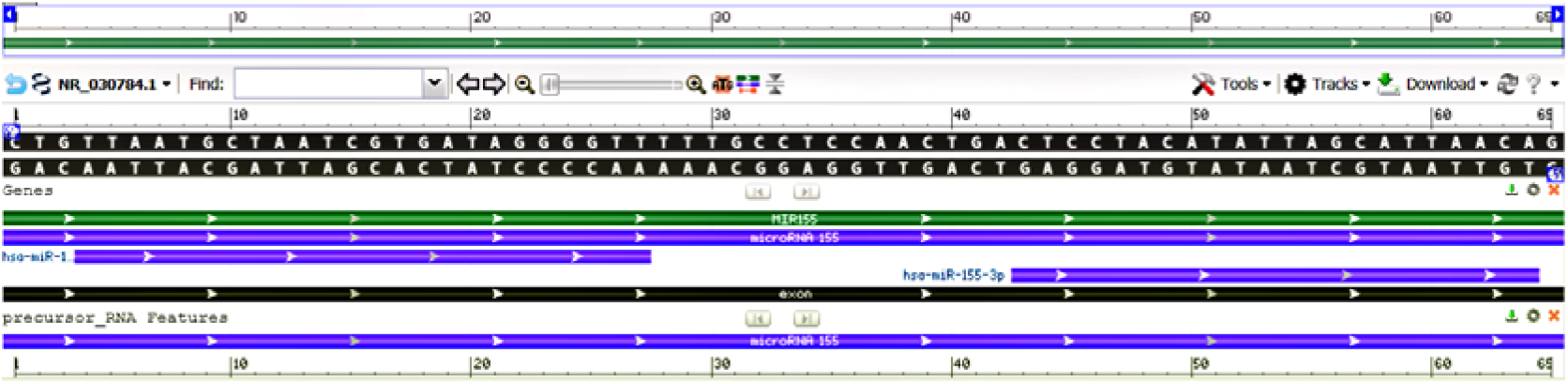
Homo sapiens microRNA 155 (MIR155) - *model-template aligment*. **Source**: https://www.ncbi.nlm.nih.gov/nuccore/NR_030784.1?report=graph

### Molecular model of microRNA 155

Based on the sequence alignment between the Homo sapiens microRNA 155 nucleotide and the template structure, the structural template for microRNA 155 was produced. Assessment tools were used para measure the reliability of the designed structure. Thereby, using the RNAstructure server comparative nucleotide modeling server, we generate a homology model of the Homo sapiens microRNA 155 (Figure 4).

**Figure 4.**
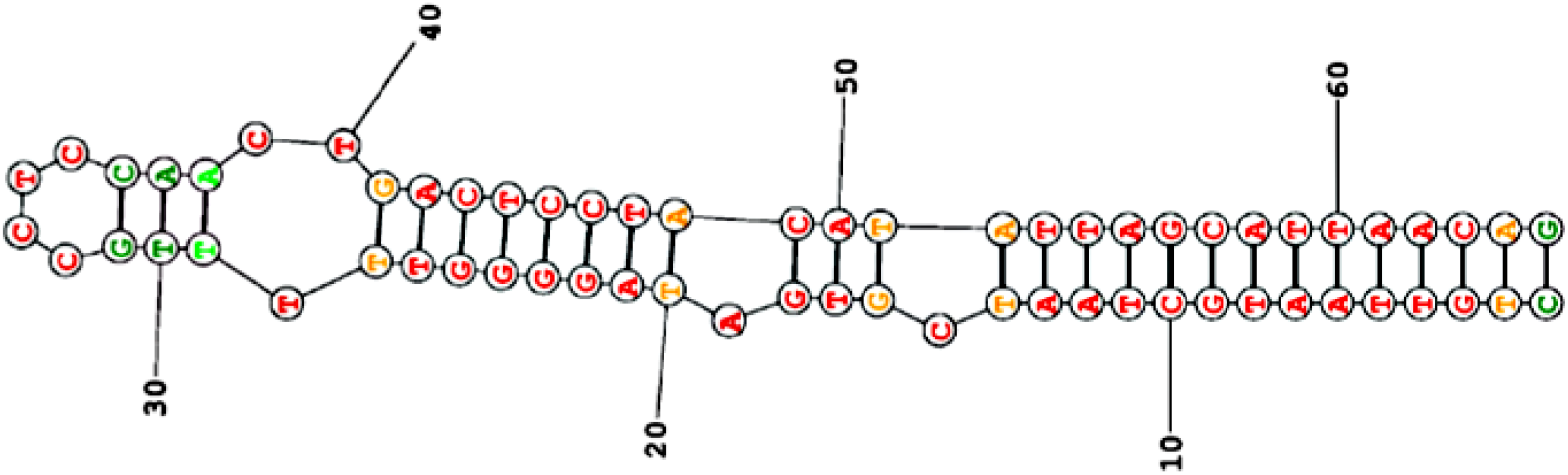
Homology model of the Homo sapiens microRNA 155 (MIR155), microRNA **Source**:https://rna.urmc.rochester.edu/RNAstructureWeb/Servers/Predict1/ResultsPages/20210703.130601-73fa4fe3/Results.html

### Nucleotide sequence of miRNA-132

To construct the structure of miRNA-132 the nucleotide sequences with the NCBI identifier code: NR_029674.1 were used under the FASTA format obtained from GenBank. The miRNA-132 was planned to encode a 101 bp linear ncRNA. Homo sapiens microRNA 132 (MIR132) analysis is demonstrated in Figure 5.

**Figure 5.**
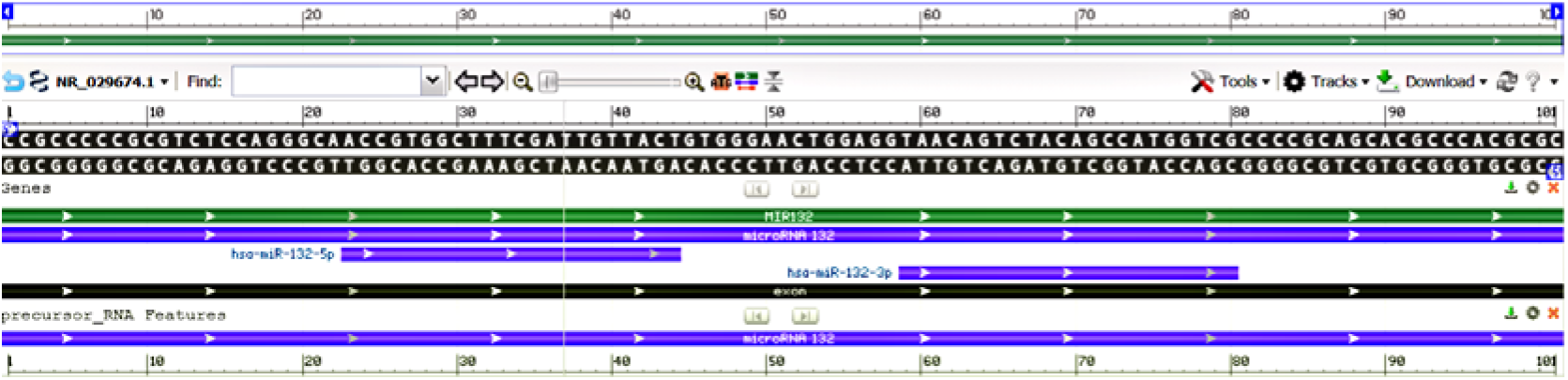
Homo sapiens microRNA 132 (MIR132) - *model-template aligment*. **Source**: https://www.ncbi.nlm.nih.gov/nuccore/NR_029674.1?report=graph

### Molecular model of microRNA 132

Nucleotide sequences of Homo sapiens microRNA 132 (MIR132) were acquired employing FASTA format; modeling was performed employing the RNAstructure server, optimized and adjusted for alignment between structural templates and microRNA 132 nucleotide. Based on sequence alignment between the template structure and microRNA nucleotide, a structural model was built for the nucleotide in question. So, employing RNAstructure server of comparative nucleotide modeling we generated a homology model of Homo sapiens microRNA 132 (MIR132), demonstrated in Figure 6.

**Figure 6.**
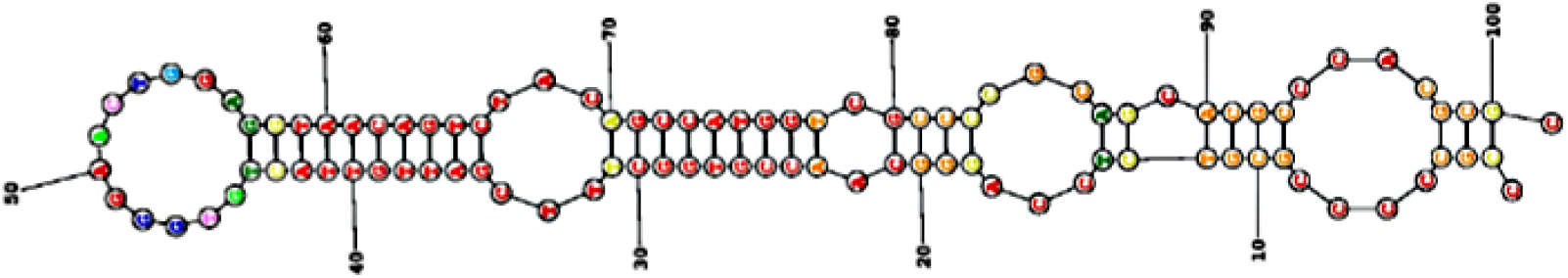
Homology model of the Homo sapiens microRNA 132 (MIR132), microRNA **Source**:https://rna.urmc.rochester.edu/RNAstructureWeb/Servers/Predict1/ResultsPages/20210703.141702-75a99147/Results.html

### Nucleotide sequence of miRNA-132

The reconstruction of miRNA-132 was performed from a nucleotide sequence archive in FASTA format obtained in GenBank database with the identifier code NCBI Reference Sequence: NR_029690.1. The miRNA-132 was planned to encode a 92 bp linear ncRNA. All coded sequences selected in FASTA format, used the annotation of the NCBI - Graphics - Homo sapiens microRNA 191 (MIR191). The microRNA-132 analysis is shown in Figure 7.

**Figure 7.**
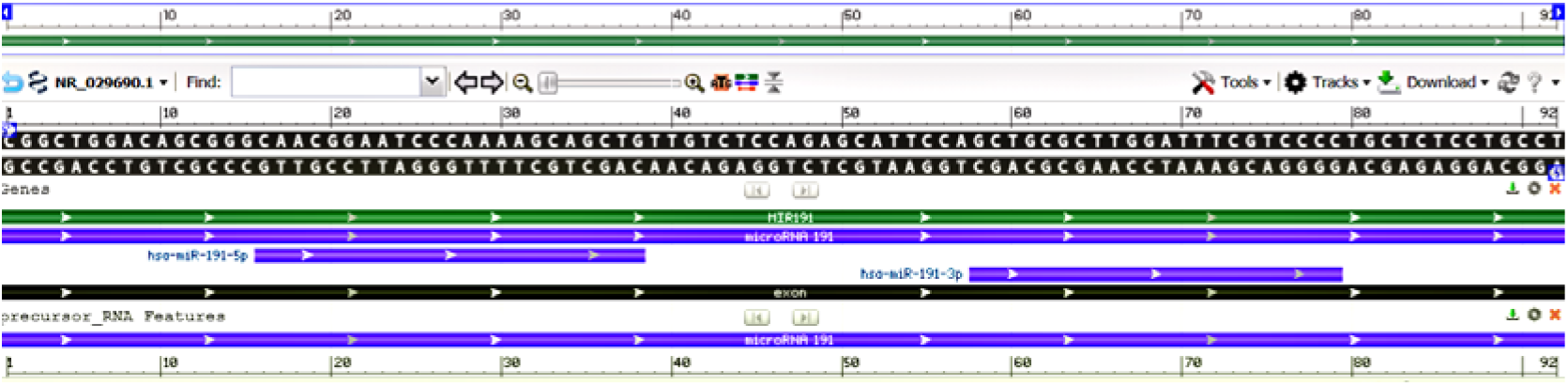
Homo sapiens microRNA 191 (MIR191) - *model-template aligment*. **Source**: https://www.ncbi.nlm.nih.gov/nuccore/NR_029690.1?report=graph

### Molecular model of microRNA-191

Based on the sequence alignment between the Homo sapiens microRNA 191 nucleotide and the template structure, the structural template for microRNA-191 was produced. Assessment tools were used para measure the reliability of the designed structure. Thereby, using the RNAstructure server comparative nucleotide modeling server, we generate a homology model of the Homo sapiens microRNA 191 (Figure 8).

**Figure 8.**
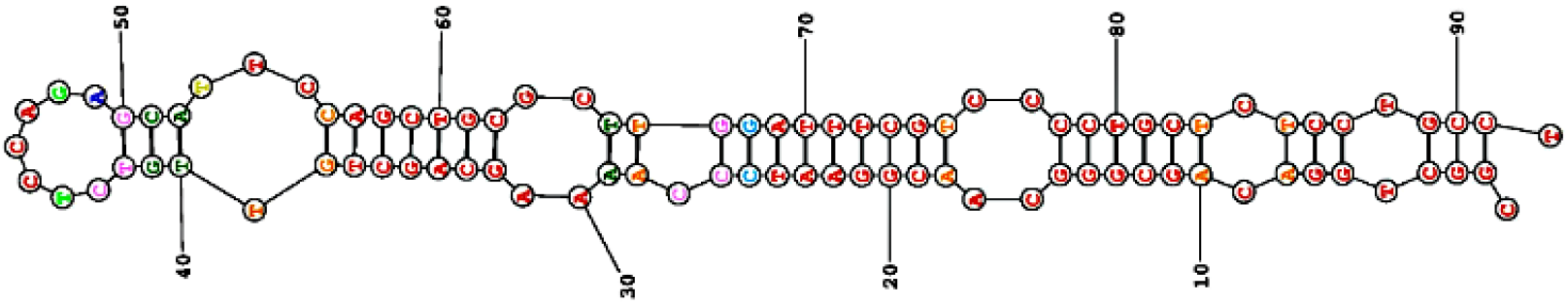
Homology model of the Homo sapiens microRNA 191 (MIR191), microRNA **Source**:https://rna.urmc.rochester.edu/RNAstructureWeb/Servers/Predict1/ResultsPages/20210703.145900-53f58714/Results.html

### Nucleotide sequence of miRNA-21

To construct the structure of miRNA-21 the nucleotide sequences with the NCBI identifier code: NR_029493.1 were used under the FASTA format obtained from GenBank. The miRNA-21 was planned to encode a 72 bp linear ncRNA. Homo sapiens microRNA 21 (MIR21) analysis is demonstrated in Figure 9.

**Figure 9.**
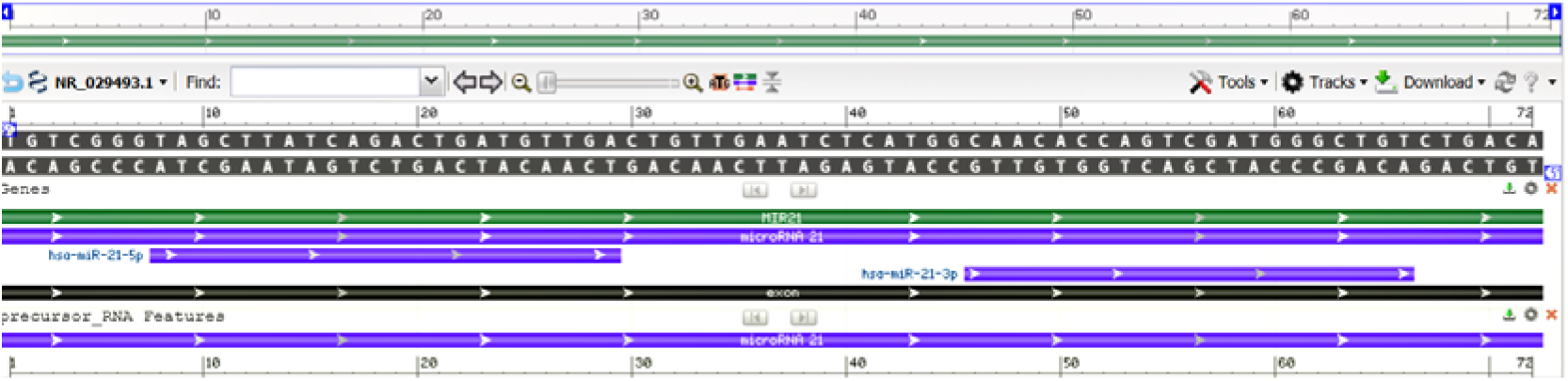
Homo sapiens microRNA 21 (MIR21) - *model-template aligment*. **Source**: https://www.ncbi.nlm.nih.gov/nuccore/NR_029493.1?report=graph

### Molecular model of miRNA-21

Nucleotide sequences of Homo sapiens microRNA 21 (MIR21) were acquired employing FASTA format; modeling was performed employing the RNAstructure server, optimized and adjusted for alignment between structural templates and miRNA-146a nucleotide. Based on sequence alignment between the template structure and miRNA-21 nucleotide, a structural model was built for the nucleotide in question. So, employing RNAstructure server of comparative nucleotide modeling we generated a homology model of microRNA 21 (MIR21), demonstrated in Figure 10.

**Figure 10.**
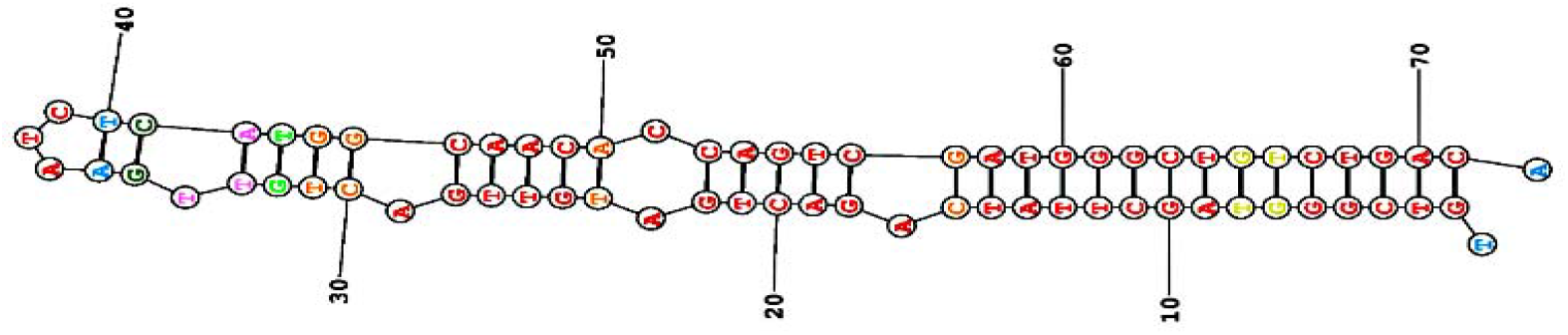
Homology model of the Homo sapiens microRNA 21 (MIR21), microRNA **Source**:https://rna.urmc.rochester.edu/RNAstructureWeb/Servers/Predict1/ResultsPages/20210708.124716-422af542/Results.html

### Nucleotide sequence of miRNA-203a

The reconstruction of miRNA-203a was performed from a nucleotide sequence archive in FASTA format obtained in GenBank database with the identifier code NCBI Reference Sequence: NR_029620.1. The miRNA-203a was planned to encode a 110 bp linear ncRNA. All coded sequences selected in FASTA format, used the annotation of the NCBI - Graphics. The Homo sapiens microRNA 203a (MIR203A), microRNA analysis is shown in Figure 11.

**Figure 11.**
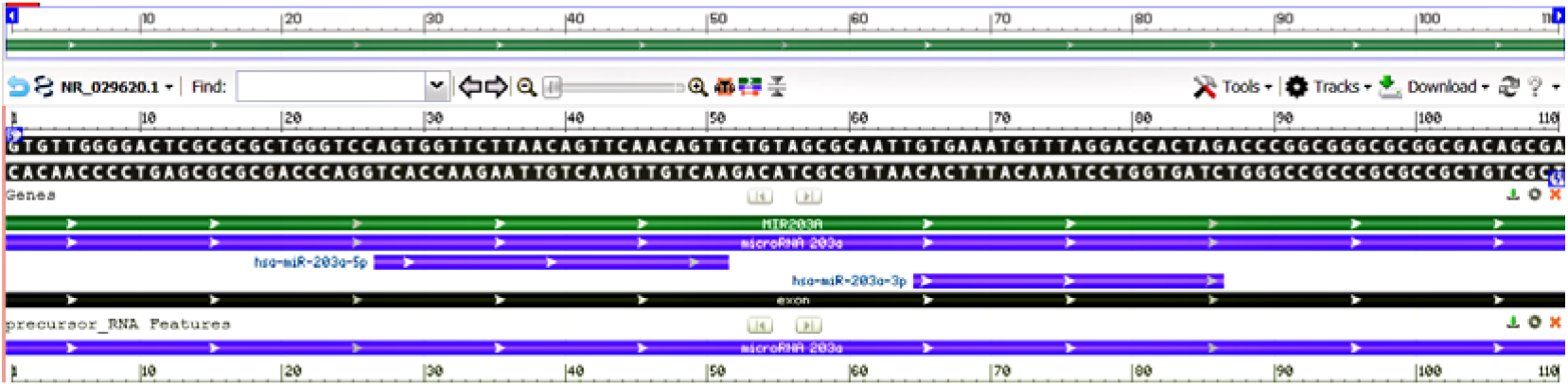
Homo sapiens microRNA 203a (MIR203A) - *model-template aligment*. **Source**: https://www.ncbi.nlm.nih.gov/nuccore/NR_029620.1?report=graph

### Molecular model of microRNA-203a

Based on the sequence alignment between the Homo sapiens microRNA 203a nucleotide and the template structure, the structural template for microRNA 203a was produced. Assessment tools were used para measure the reliability of the designed structure. Thereby, using the RNAstructure server comparative nucleotide modeling server, we generate a homology model of the Homo sapiens microRNA 203a (MIR203A) (Figure 12).

**Figure 12.**
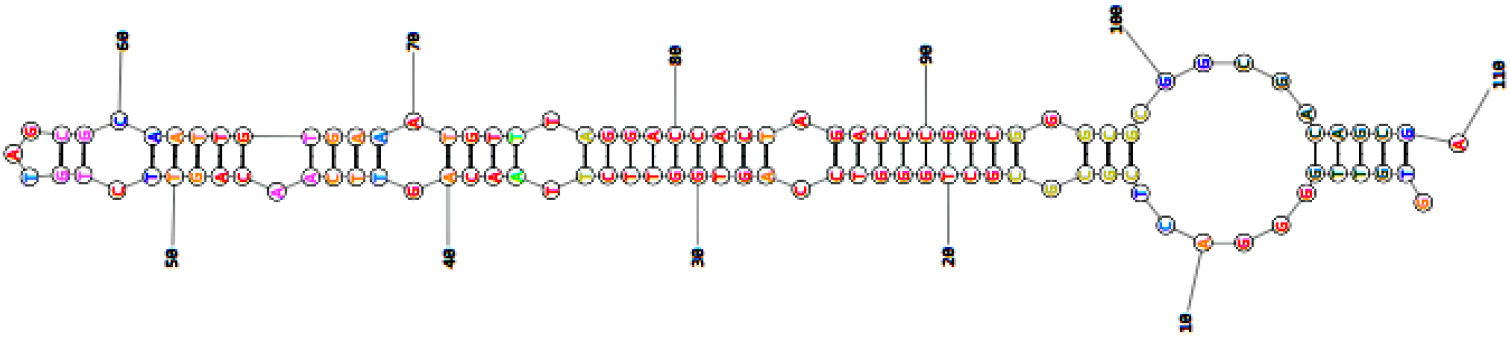
Homology model of the Homo sapiens microRNA 203a (MIR203A). **Source**:https://rna.urmc.rochester.edu/RNAstructureWeb/Servers/Predict1/ResultsPages/20210708.132036-491e2e54/Results.html

### Nucleotide sequence of miRNA-203b

To construct the structure of miRNA-203B the nucleotide sequences with the NCBI identifier code: NR_039859.1 were used under the FASTA format obtained from GenBank. The miRNA-146B was planned to encode a 86 bp linear ncRNA. Homo sapiens microRNA 203b (MIR203B), microRNA analysis is demonstrated in Figure 13.

**Figure 13.**
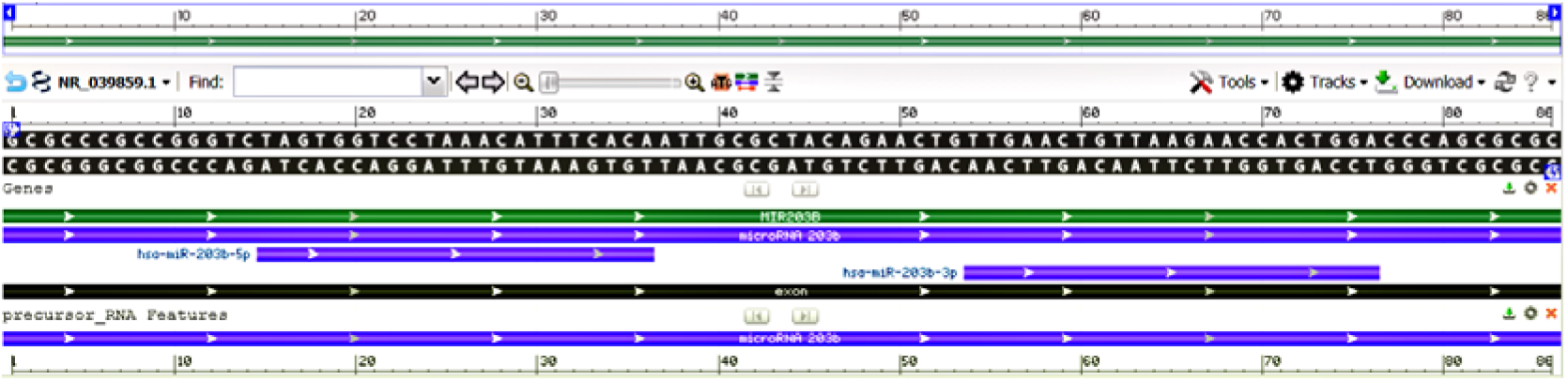
Homo sapiens microRNA 203b (MIR203B) - *model-template aligment*. **Source**: https://www.ncbi.nlm.nih.gov/nuccore/NR_039859.1?report=graph

### Molecular model of microRNA-203b

Nucleotide sequences of Homo sapiens microRNA 203b (MIR203B), microRNA were acquired employing FASTA format; modeling was performed employing the RNAstructure server, optimized and adjusted for alignment between structural templates and miRNA-203b nucleotide. Based on sequence alignment between the template structure and miRNA-203b nucleotide, a structural model was built for the nucleotide in question. So, employing RNAstructure server of comparative nucleotide modeling we generated a homology model of microRNA 203b (MIR203B), demonstrated in Figure 14.

**Figure 14.**
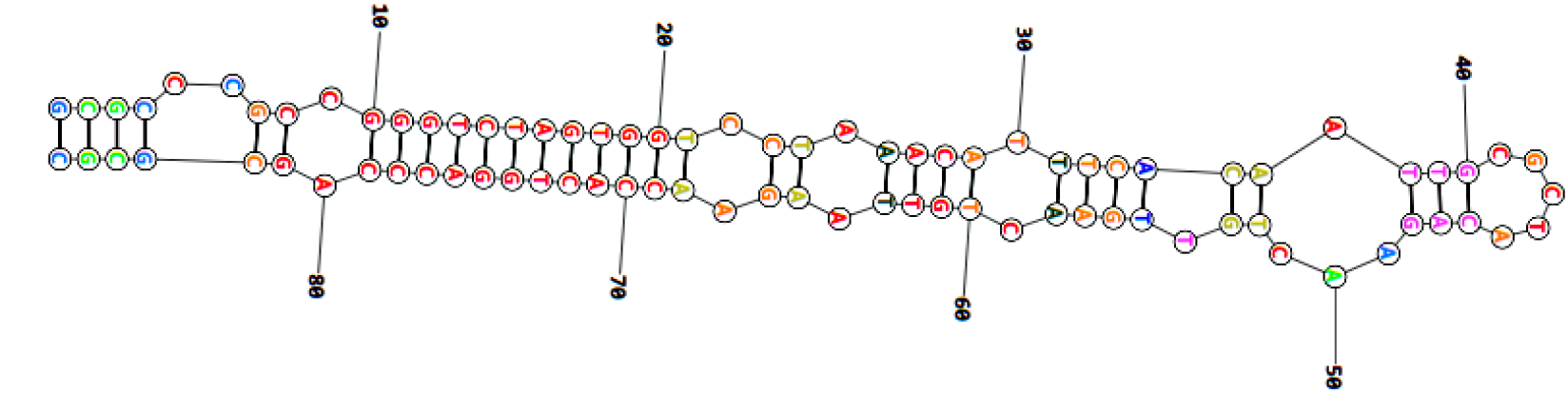
Homology model of the Homo sapiens microRNA 203b (MIR203B), microRNA **Source**:https://rna.urmc.rochester.edu/RNAstructureWeb/Servers/Predict1/ResultsPages/20210708.140010-4b987349/Results.html

### Nucleotide sequence of miRNA-210

The reconstruction of miRNA-155 was performed from a nucleotide sequence archive in FASTA format obtained in GenBank database with the identifier code NCBI Reference Sequence: NR_029623.1. The miRNA-210 was planned to encode a 110 bp linear ncRNA. All coded sequences selected in FASTA format, used the annotation of the NCBI - Graphics. The Homo sapiens microRNA 210 (MIR210) analysis is shown in Figure 15.

**Figure 15.**
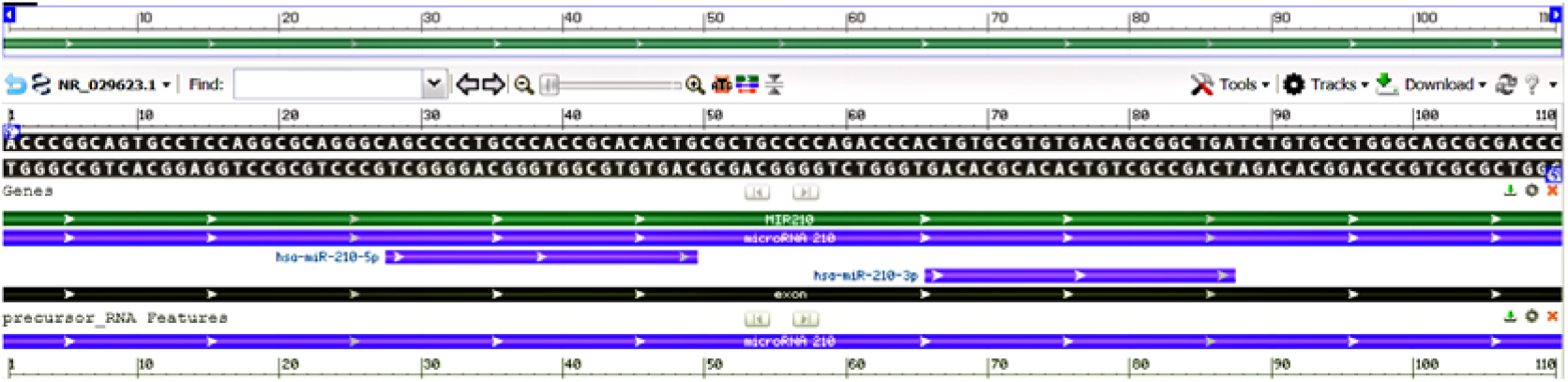
Homo sapiens microRNA 210 (MIR210) - *model-template aligment*. **Source**: https://www.ncbi.nlm.nih.gov/nuccore/NR_029623.1?report=graph

### Molecular model of microRNA-210

Based on the sequence alignment between the Homo sapiens microRNA-210 nucleotide and the template structure, the structural template for microRNA-210 was produced. Assessment tools were used para measure the reliability of the designed structure. Thereby, using the RNAstructure server comparative nucleotide modeling server, we generate a homology model of the Homo sapiens microRNA-210 (Figure 16).

**Figure 16.**
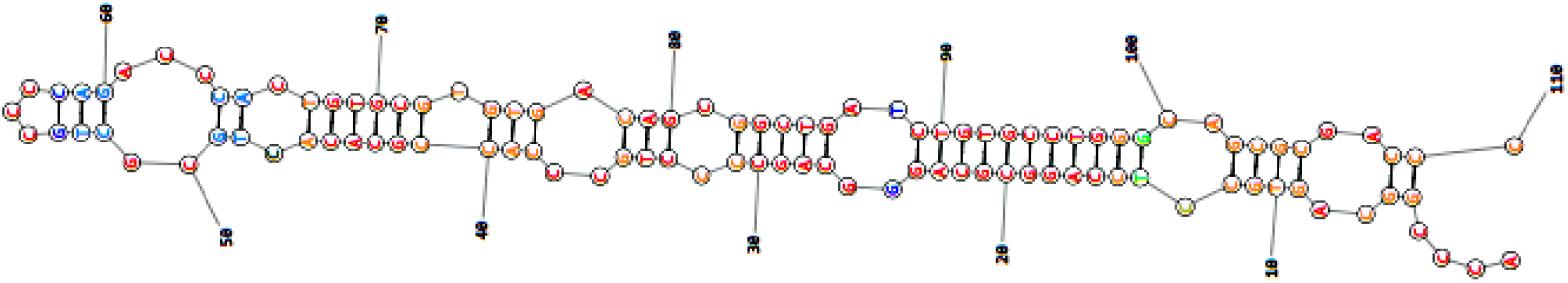
Homology model of the Homo sapiens microRNA 210 (MIR210), microRNA **Source**:https://rna.urmc.rochester.edu/RNAstructureWeb/Servers/Predict1/ResultsPages/20210708.143619-218c16fb/Results.html

## DISCUSSION

The miRNAs are small non-protein-coding RNA which has the controlling role in multiples physiological and pathophysiological functions. About a decade ago the knowledge about the structure and performance of miRNAs has increased significantly. Our study presents a tutorial on molecular modeling and demonstrates in silico the projection of the molecular structure of 8 miRNAs overexpressed in diabetic foot ulcers and projecting *in silico* their molecular structures.

Diabetic foot ulcers are serious complications of diabetes mellitus contributing to a high overall of morbidity and mortality, and it is necessary to understand its genetic and molecular basis to increase efficiency of its treatment. Various microRNAs and their target genes participate to tissue fibrosis of diabetic foot ulcers, in the same way that several microRNAs participate of angiogenesis which is committed in the vascular tissue of the diabetic individual.^6,7^

The miRNA-146a is express in chromosome 5 with location 5q33.3, and is implicated in pathogenesis of many diseases by inhibiting the manifestation of its targets. The miRNA-146 family is composed of the miRNA-146a and miRNA-146b that regulate negatively the cellular inflammatory gene expression, including monocytes, endothelial cells, and epithelial cells, moreover, the miRNA-146a induces innate immune response upon contact with antigens.^8^ The miRNA-146a affects the inflammatory response and its expression is significantly reduced in the diabetic foot ulcers. However, the treatment of diabetic foot ulcers presented increased expression of miRNA-146a in the around tissue, demonstrating that the expression of miRNA-146a can be dependent on an extracellular signal.^9^ Furthermore, there is a study showing that treatment with miR-146a inhibitor is helpful in speed wound healing in diabetic foot ulcers in function of increased vascular endothelial growth factor and fibronectin production.^10^

The post-genomic era has featured technological developments favoring the expansion of microarrays and databases, but the big hurdle for researchers is to exploit the information that is useful to them. Bioinformatics has several tools to predict the structure of microRNAs, leading to an understanding of the mode of transcriptional patterning of microRNAs. We explore the miRNA-146a sequences in the NCBI database through of GenBank database based in identifier NCBI, distinguishing all the nucleotide encoded in miRNA-146a and designing their structure by applying domain analysis tools.

Bioinformatics tools for miRNA prediction have won accessibility since experimental studies to set miRNAs are uncommon in their realization. The in silico evaluation of miRNAs is reasoned mainly on primary and secondary structure performance analysis. Analyzing the literature, we found no study with projection of miRNA-146a with two-dimensional (2-D) structure model. For our analysis, we evaluated nucleotide database GenBank, and a 2-D model of miRNA-146a was designed with the online RNAstructure program.

The miRNA-155 is a small, non-coding RNA, single-stranded originated and identified in human chromosome 21 and with location 21q21.3, previously known as B-cell Integration Cluster. MiRNA-155 in humans is codified by the MIR155 host gene or MIR155HG.^11^ MiRNA-155 is involved in various physiological and pathological processes. Studies have demonstrated the possible implication of miR-155 in pathogenesis of diabetic complications.^12^

Treating a chronic diabetic foot ulcer is a difficult procedure because the disability to at the same time treats the inflammatory process and the re-epithelization, because both increase bacterial colonization increase bacterial colonization delaying ulcer healing. The miR-155 has been associated increase the severity of diabetic foot ulcers.^4^ Diabetic subjects they have reduced levels of miRNA-155 expression in mononuclear cells and peripheral, and the miRNA-155 inhibition leads to a decrease in the inflammatory process, due to the increase of M2 macrophage polarization and increase in type I collagen deposition.^13^ In addition, the miR-155 in immune cells plays a pro-inflammatory role by means of suppression of cytotoxic T-lymphocyte-associated protein 4.^14^

In our literature review, we found only one study with a 2-D structural model of miRNA-155.^15^ In our study, the sequenced analysis of the miRNA-155 was assessed and designed in the 2-D structural model. The nucleotide analysis of miRNA-155 was realized through of FASTA format, and for 2D modeling we use the RNAstructure program, tuning and optimizing for alignment between structural templates and miRNA-155. So, to build the 2-D structural model of miRNA-155 we used structural homology analysis strategy with the Nucleotide database sequence. The use of molecular models should promote progress the development of new drugs facilitating therapy in target organs.

The miRNA-132 is transcribed in an intergenic region on chromosome 17 in humans, and with location 17p13.3.^16^ The miRNA-132 have various targets including mediators of angiogenesis and inflammation, that are fundamental in diabetic foot ulcer. miRNA-132 can lead to multiplication of endothelial cells and has been involved in neovascularization through angiogenic factors.^17^ Conversely, miRNA-132 can also modulate inflammation, promoting inflammation that at the base of insulin resistance of the diabetes mellitus.^18^ Experimental study wound healing showed the efficacy of miRNA-132 in diabetic mouse model, and a case study in diabetic human employing topical application of miRNA-132 formulated also demonstrated that it was efficient to treat the diabetic wound.^19^ Revising the medical literature, we did not find any scientific study showing the 2-D projection structure of the miRNA-132. We design the 2-D molecular model of miRNA-132 making use of the homology modeling method reasoned on the miRNA-132 sequence and the high-homology structure as the template using RNAstructure web server based on the nucleotide sequence of miRNA-132 of NCBI-Nucleotide database.

The miRNA-191 is a member of the miR-191 family miR-191 located on human chromosome 3 (3p21.31 position).^20^ The modulation in miR-191 expression control the cell proliferation, apoptosis, and cell cycle in several diseases, furthermore, miR-191 is a stress-sensitive miRNA and low levels of this miRNA have been related to diabetes mellitus.^21^ A recent study has shown that plasma miRNA-191 profiles presents a unidirectional changes in circulating plasma levels from diabetics patients with impaired wound healing compared with diabetics with no chronic foot ulcer.^22^ We conducted a literature review and did not find any manuscripts secondary structure model of the miRNA-191. We model computationally in 2-D structure the Homo sapiens miRNA-191, using the modeling method by homology based on the FASTA sequence of miRNA-191 and a high-homology structure with the RNAstructure web server was built.

The human microRNA-21 is located in chromosome 17q23.2 within a coding gene called vacuole membrane protein, and performs a key role in many diseases, as well as in biological functions developmental and regulations of several processes immunological.^23^ It has been shown that that miR-21 expression presents down-regulated in the course of diabetic wound healing, being crucially important in fibroblast migration, collagen deposition, as well as presents pro-proliferative, anti-apoptotic, and anti-inflammatory properties.^24^ In diabetic wound, the miRNA-21 performs role at the different stages of wound healing process, presenting high in the inflammatory phase, reducing in proliferative phase, and significantly elevating in remodeling phase.^25^ In our literature review, we found only one study with a 2-D structural model of miRNA-21.^26^ We design the 2-D molecular model of miRNA-21 making use of the nucleotide database source GenBank based on the FASTA sequence of miRNA-21 and designed with the online RNAstructure server.

The microRNA-203a and microRNA 203b Homo sapiens (human) are located in chromosome 14q32.33. The microRNA-203 modulates inflammatory reaction, play important role in cell proliferation, being of great importance in the development of the skin.^27^ In ulcers of diabetic individuals increased expression levels of miRNA-203, which is a miRNA highly abundant in epidermis and keratinocyte-specific miRNA that is highly abundant in epidermis, and miRNA-203 can play a significant role in diabetic wound healing, and was shown to be positively correlated with diabetic foot ulcer severity.^28^ Analyzing the literature, we found no study with projection of miRNA-203 with 2-D structure model. In our analysis, we evaluated nucleotide database GenBank, and a 2-D model of miRNA-203a and miRNA-203b was projected with the online RNAstructure program.

The miRNA-210 is express in chromosome 11, and with location 11p15.5.The miR-210 is implicated in several biological processes of human body, and regulating various cellular functions as metabolism, cell proliferation, apoptosis, and angiogenesis.^29^ In diabetic foot ulcers the miR-210 expression is reduced, and local reconstitution using mimicry miR-210 heals diabetic wound significantly, in function of decrease in the oxygen consumption rate through the wound due to the local reduction of Reactive Oxygen Species levels in the wound tissue.^30^ Reviewing the medical literature, we observed that none study with secondary structure model of the miRNA-210 was published. We used data of FASTA sequence to build the model computationally of 2-D structure the Homo sapiens miRNA-210, and a high-homology structure with the RNAstructure server.

## CONCLUSION

The structure and function of miRNAs are established by nucleotides sequences and structure prediction methods allow identifying the location of binding sites on nucleotides of essential value for use clinical and pharmacological.

We show *in silico* secondary structures design of selected of 8 miRNAs overexpressed in diabetic foot ulcers healing by means of computational biology.

